# Noisy circumnutations facilitate self-organized shade avoidance in sunflowers

**DOI:** 10.1101/2022.06.11.495747

**Authors:** Chantal Nguyen, Imri Dromi, Aharon Kempinski, Gabriella E. C. Gall, Orit Peleg, Yasmine Meroz

## Abstract

Circumnutations are widespread in plants and typically associated with exploratory movements, however a quantitative understanding of their role remains elusive. In this study we report, for the first time, the role of noisy circumnutations in facilitating an optimal growth pattern within a crowded group of mutually shading plants. We revisit the problem of self-organization observed for sunflowers, mediated by shade response interactions. Our analysis reveals that circumnutation movements conform to a bounded random walk characterized by a remarkably broad distribution of velocities, covering three orders of magnitude. In motile animal systems such wide distributions of movement velocities are frequently identified with enhancement of behavioral processes, suggesting that circumnutations may serve as a source of functional noise. To test our hypothesis, we developed a Langevin-type parsimonious model of interacting growing disks, informed by experiments, successfully capturing the characteristic dynamics of individual and multiple interacting plants. Employing our simulation framework we examine the role of circumnutations in the system, and find that the observed breadth of the velocity distribution represents a sharp transition in the force-noise ratio, conferring advantageous effects by facilitating exploration of potential configurations, leading to an optimized arrangement with minimal shading. These findings represent the first report of functional noise in plant movements, and establishes a theoretical foundation for investigating how plants navigate their environment by employing computational processes such as task-oriented processes, optimization, and active sensing. Since plants move by growing, space and time are coupled, and dynamics of self-organization lead to emergent 3D patterns. As such this system provides conceptual insight for other interacting growth-driven systems such fungal hyphae, neurons and self-growing robots, as well as active matter systems where agents interact with past trajectories of their counterparts, such as stigmergy in social insects. This foundational insight has implications in statistical physics, ecological dynamics, agriculture, and even swarm robotics.

**One sentence summary of paper:** The study highlights noisy circumnutations as a strategy plants use for optimizing growth in crowded conditions.

## Introduction

The survival of plants greatly depends on light availability. In many natural habitats, multiple neighboring plants shade each other, competing over this critical resource. The presence of neighbors varies over space and time, and plants have evolved the ability to detect neighbors and respond by adapting their morphology accordingly [1, 2, 3, 4, 5]. Indeed, a fundamental difference between plants and other motile organisms is that plants generally move by growing; an irreversible process which imbues plant movement with a commitment to the permanent morphology. The direction of plant growth is dictated either by external directional cues such as light, processes termed tropisms, or by inherent internal cues, such as the exploratory periodic movements termed *circumnutations*. Recently, Pereira et al. [6] found that sunflower crops growing in a row at high densities self-organized into a zigzag conformation of alternating inclined stems, thus collectively increasing light exposure and seed production. Self-organized processes refer to initially disordered systems where order arises from local interactions between individuals, facilitated by random perturbations, or noise. Local interactions were found to be mediated by the shade avoidance response, a form of tropism where plant organs grow away from neighboring plants [7], responding to changes in the ratio between red and far-red wavelengths, characteristic of the light spectrum of plant shade [8, 9, 10]. However, the source of perturbations required for the observed self-organization in this system remains elusive.

Noise plays a critical role in self-organized systems: at the right magnitude relative to the interactions, it enables the system to explore a variety of states, thus enabling reliable adaptation to short-term changes in the environment while maintaining a generally stable behavior [11, 12, 13, 14]. Too little noise confines the system to a sub-optimal state, while too much noise masks the interactions. In biological systems, noise is often identified as being *functional*, exhibiting a wide spectrum of manifestations. For example, organisms use noise in order to increase sensory salience, ultimately enabling them to balance the behavioral conflict between producing costly movements for gathering information (“explore”) versus using previously acquired information to achieve a goal (“exploit”) [15, 16, 17]. Additionally, the navigation paths of bacteria [18], insects [19], and mammals [20, 21] exemplify the intricate trajectories adopted to counterbalance uncertain surroundings [22]. Likewise, the behavioral variability observed in honeybees, specifically in the triggering of fanning, serves as a mechanism for ventilating their hives [23]. In all of these instances, noisy processes play a fundamental role in facilitating biological functionality.

Here we propose cirumnutations as the predominant source of perturbations in this system. These movements are ubiquitous in plants, following elliptical or irregular trajectories with large variations in both amplitude and periodicity [24, 25, 26, 27, 28]. Circumnutations assist climbing plants in locating mechanical supports [24, 29], facilitate root navigation around obstacles [30, 31], and contribute to the regulation of shoot stability during elongation growth and tropic bending [32, 33]. However, their ecological function in non-climbing shoots, which constitute the majority of plants, remains unclear [34, 35, 28, 36].

To test our hypothesis that noisy circumnutations can benefit collective plant growth, we record movements of mutually shading sunflowers in a controlled environment, and develop a Langevin-type minimal model where mutually shading sunflower crowns are represented by interacting growing disks, and the quasi-periodic circumnutation movements are approximated by random perturbations with equivalent statistics. The model, informed and validated by our experimental data, recovers the observed dynamics of sunflower growth patterns, and enables us to examine the functionality of circumnutations.

Our model finds that the characteristic statistics of measured circumnutations are such that they maximize the light exposure of the system, suggesting that circumnutations play a critical role in reaching collectively optimal growth configurations under light competition.

## Results

### Recapitulating alternate inclination experiments

We recapitulate the self-organized zigzag growth conformation observed by Pereira *et al*. [6] in a controlled environment, as described schematically in Fig. 1C, enabling us to record the crown dynamics throughout. The slow dynamics of the sun was shown not to affect the observed staggered formation [6], and therefore a fixed light source provides a good approximation for field experiments. Fig. 1A shows an initial arrangement of five young sunflowers (*Helianthus annuus*) approximately 7 days old, where the plant crowns and their centers are highlighted, clearly aligned to a horizontal line. We define the position of crown i at time *t* as **r**_i_(*t*), and define the average center-to-center distance between adjacent plants *d*_CC_(*t*) (defined in Eq. 8 in the Methods), normalized by the initial center-center distance at *t* = 0 so that *d*_CC_(0) = 1 by definition. This value serves as a measure of how closely (*d*_CC_ *<* 1) or sparsely (*d*_CC_ *>* 1) the plant crowns are distributed. We allow the plants to grow undisturbed over 7 days, following the position and size of the crowns throughout. Fig. 1B shows their final configuration, where the crowns have grown in size, and their centers are clearly in a staggered formation. In this particular example, the average distance has increased such that *d*_CC_ = 1.2, ascribed to the zigzag configuration. A time-lapse is brought in Video 1 in the Supplementary Material (SM). The trajectories of the tracked crown centers over the course of this specific multiple-plant experiment are shown in Fig. 1D. Over 12 such experiments, we observe that the final center-to-center distance is generally greater than the initial distance, with an average value of *d*_*CC*_ = 1.21 (Fig. 1E), indicating that crowns indeed deflect away from one another in dense growth setups.

**Figure 1:**
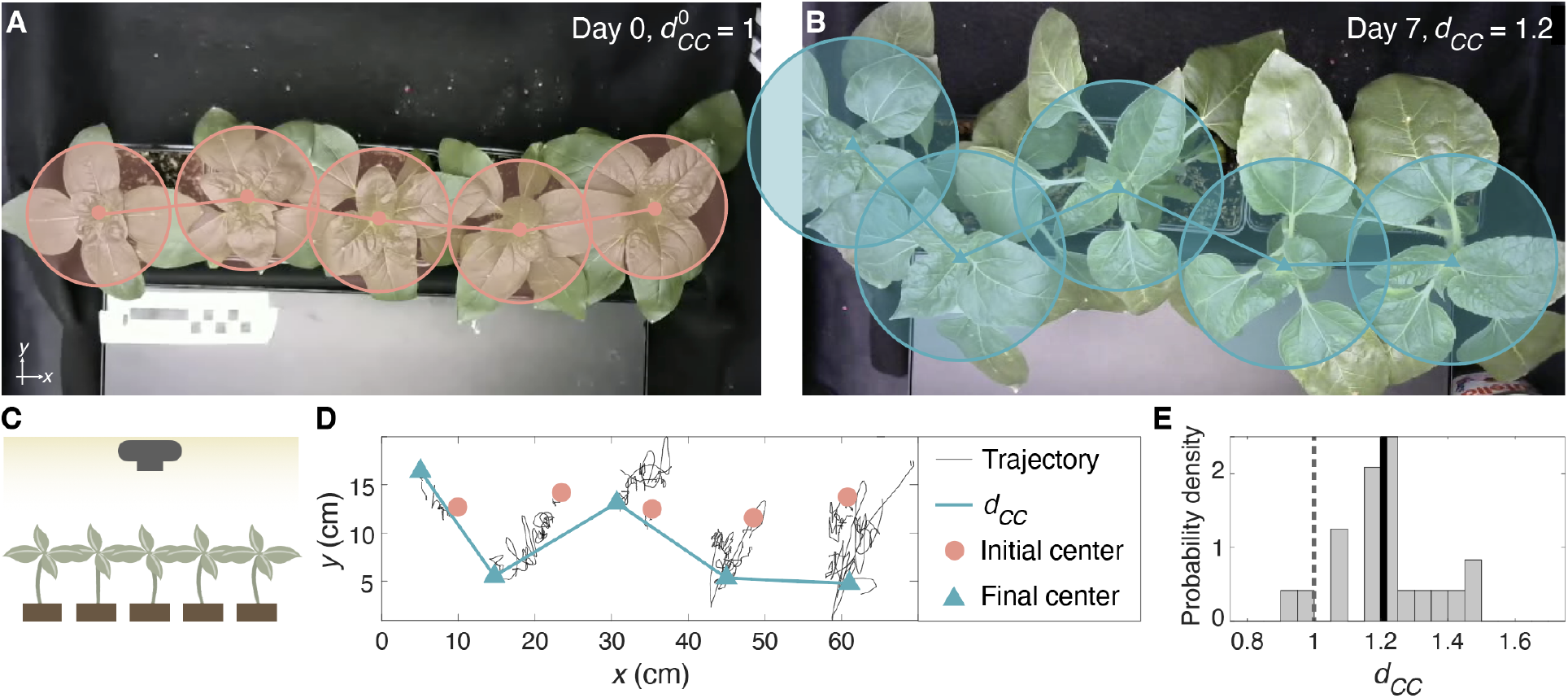
Plants grown in a dense row deflect away from each other to minimize mutual shading. **(A)** Snapshot of five sunflowers (*Helianthus annuus*) placed in a row, on the first day of recording. Pink dots indicate crown centers, and circles illustrate the model representation of crowns as disks in the *x*-*y* plane. The initial average center-to-center separation between pairs of adjacent plants, represented by connecting lines, is normalized such that 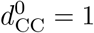 (Eq. 8). **(B)** After seven days, plants deflect from the center line, captured by the increased center-to-center distance *d*_CC_ = 1.2. Blue triangles indicate crown centers, and circles represent crowns. **(C)** Schematic of experimental setup: five plants are arranged in a row, with homogeneous lighting placed overhead, and their dynamics are recorded from above. **(D)** Trajectories of crown centers during a sample 7-day recording are shown (black lines), where initial and final crown positions are represented by pink dots and blue triangles, respectively. Blue lines illustrate the increased center-to-center separation *d*_CC_ between pairs of adjacent plants, highlighting the arising deflected pattern. **(E)** Histogram of the final *d*_CC_ values from 12 multiple-plant experiments, indicating a final separation with a mean *d*_*CC*_ = 1.21 ± 0.14 (solid line), greater than that of the initial separation *d*_CC_ = 1 (dashed line).

### Langevin-type minimal model for dynamics of noisy interacting crowns

We formulate the general framework for a minimal model describing interacting plant crowns in the 2D plane based on Langevin-type description. We approximate growing crowns with circular disks whose radius *R*_C_(*t*) increases according to a measured growth rate (Fig. 2A). As in the experiments, **r**_i_(*t*) describes the position of the crown center of plant i at time *t*. We approximate the sunflower shade avoidance response, and other possible contributions such as mechanical interactions (see Video 1 in the SM), as an effective pairwise interaction between disks. When two crowns i and j overlap, *r*_ij_(*t*) = |**r**_i_(*t*) − **r**_j_(*t*)| *<* 2*R*_C_(*t*), they experience an effective repulsive *shading force* 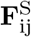 (Fig. 3A) which, in the absence of a clear relation, we assume follows a simple inverse-square relation:

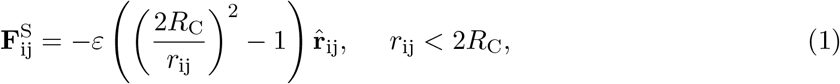

where *r*_ij_ = |**r**_ij_| = |**r**_i_ − **r**_j_| is the center-to-center distance between two crowns i and j at time *t* (explicit time dependence omitted for clarity),and 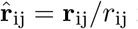 is a unit vector. The coefficient *ε* scales the magnitude of the force and is determined from experiments, as detailed in the next section. Here 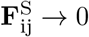 for *r*_ij_ → 2*R*_C_ avoiding a jump discontinuity at *r*_ij_ = 2*R*_C_, and we fix 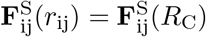 for *r*_ij_ *< R*_C_ to prevent extremely large forces in simulations. The forces are symmetric, i.e. 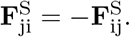. For a system of multiple mutually shading crowns, the force acting on crown i is the sum of all pairwise forces with other crowns 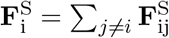 (Fig. 3A). In the SM we show that the resulting dynamics are robust to different powers in Eq. 1 (Fig. S2). We represent the perturbations driven by circumnutation movements with a noise term ***η***, where we assume step sizes *η* are taken from a distribution *P* (*η*) in random directions such that ⟨***η***⟩ = 0, and uncorrelated in time such that ⟨*η*(*t*)*η*(*t′*)⟩ = *σ*^2^ *δ*(*t* − *t′*) where the variance *σ*^2^ is given by:

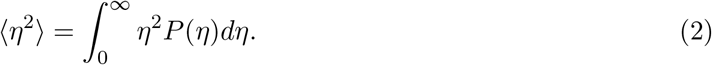

**Figure 2:**
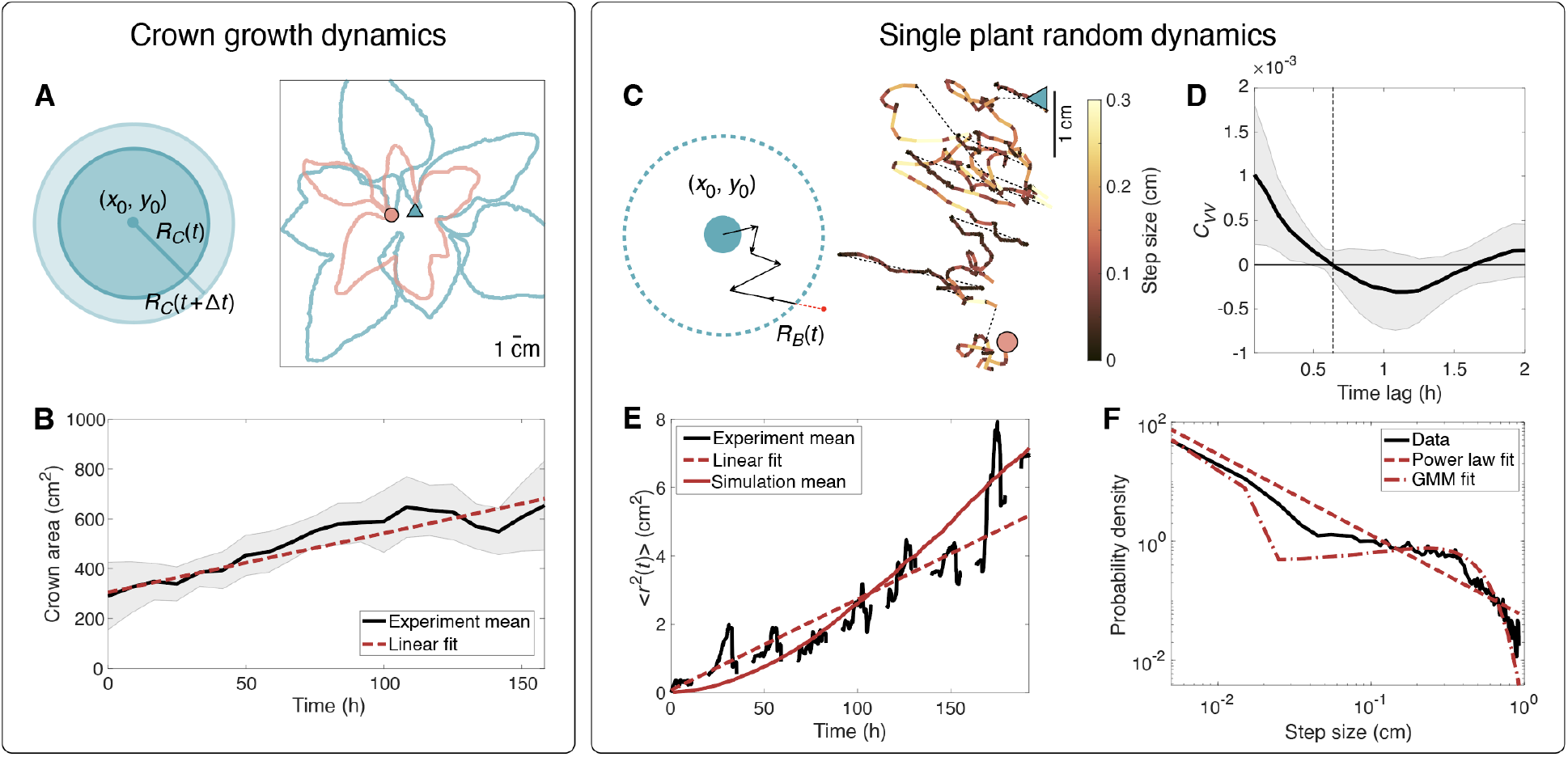
Characterization of single-plant dynamics. (Left) Crown growth dynamics. **(A)** Left: graphical representation of model; plant crown is approximated with a circle of radius *R*_C_(*t*) that increases according to the growth rate 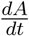. Right: example outlines of a segmented plant crown at the beginning (pink) and after 10 days (blue). **(B)** Crown area, 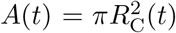, as a function of time, averaged over *N* = 8 experiments (solid black line); shaded area represents 1 standard deviation. Each experiment is individually fit to a line; the average fit (dashed red line) yields a slope 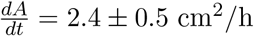 (Eq. 4). **(Right) Circumnutation dynamics (C)** Left: graphical representation of model; the crown (blue circle) is bounded by a reflective boundary of radius *R*_B_(*t*) (dashed line), increasing with time following Eq. 5. The crown follows a random walk (RW) and takes a step in a random direction at each time step; if a step crosses the boundary (red arrow) it is reflected back. Right: example of measured crown trajectory. The initial and final positions of the crown, after 10 days, are marked by a pink dot and blue triangle, respectively. Gaps in the trajectory indicate the nightly 8h “dark” period (not recorded). Color code represents local velocity (light colors represent a larger step size, and therefore higher velocity). **(D)** The autocorrelation of steps *C*_vv_ as a function of time lag shows that at a time lag of Δ*t* = 0.6 h, steps are uncorrelated. **(E)** The mean squared displacement (MSD) ⟨*r*^2^(*t*)⟩ across the length of single plant experiments, averaged over all plants (solid black line). The gaps in the curve represent 8-hour nighttime periods where the plant is not recorded. The dashed red line represents the linear fit in Eq. 5, ⟨*r*^2^(*t*)⟩ = 0.027*t* + 0.074, with *R*^2^ = 0.84. The solid red line represents the MSD averaged over 5000 simulation instances of single plants. **(F)** The step size distribution (black line). The sampling rate of trajectories is based on the autocorrelation of steps *C*_vv_ (panel (D)). The resulting distribution of step sizes (equivalent to velocities) is wide, spanning three orders of magnitude, and can be approximated to a power law (dashed red line) yielding *P* (*η*) = 0.052*η*^*−*1.40^ with *R*^2^ = 0.802 (Eq. 6), or a 2-parameter Gaussian mixture model (GMM with means 0.0060 and 0.229 with mixing proportions 0.59 and 0.41, respectively, *R*^2^ = 0.818, dash-dot red line).

**Figure 3:**
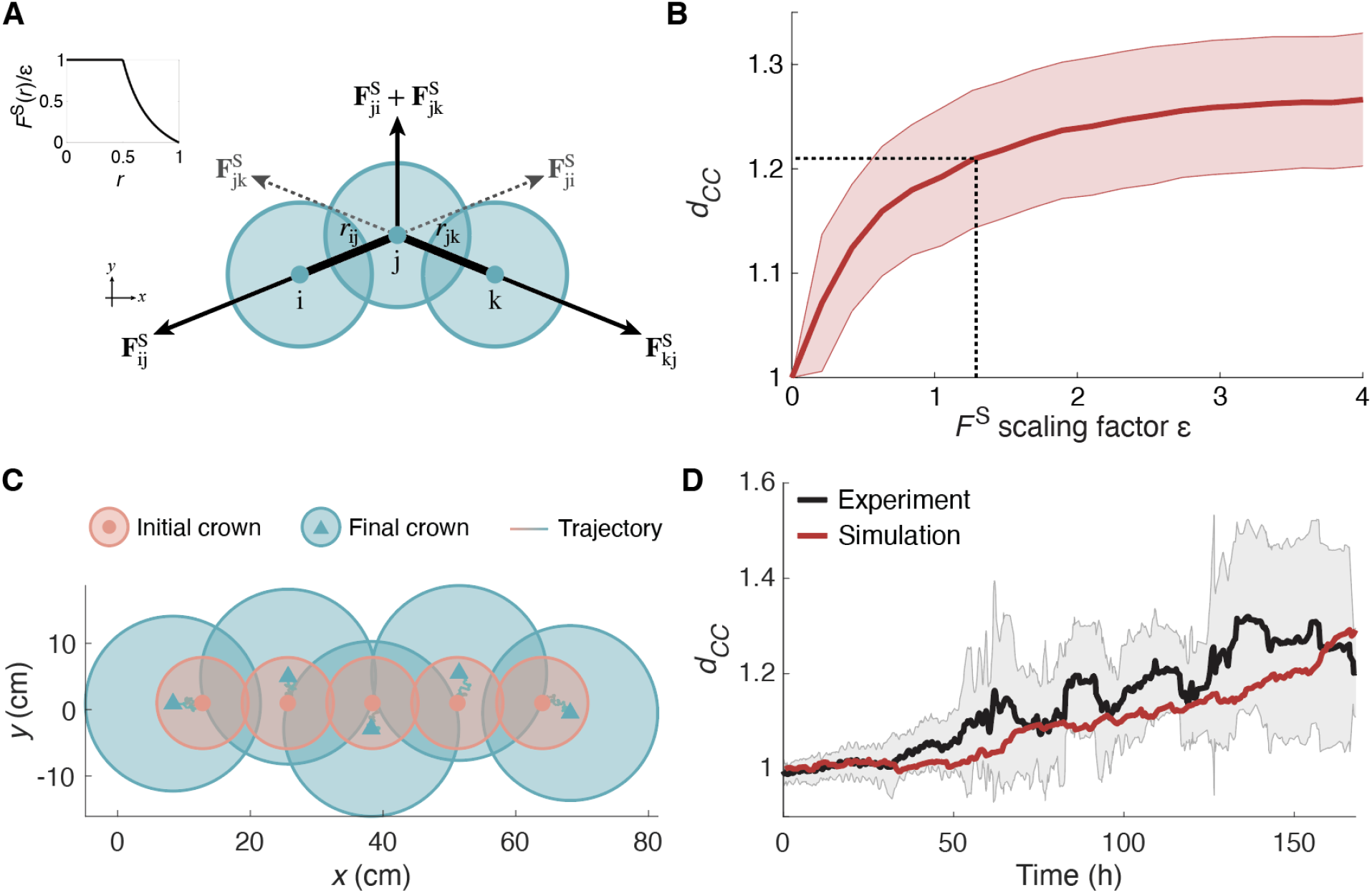
Plant-plant interaction simulations recover self-organization dynamics. **(A)** The shade response of mutually-shading crowns are represented by repulsive forces **F**^S^ (Eq. 1). As an example, three crowns are placed in a staggered formation, such that j is equidistant from i and k, experiencing repulsive forces from both: 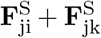. The *x*-components of these forces cancel, leading to a net force in the *y*-direction. The outer crowns i and k are affected by the central crown, with reciprocal forces 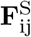 and 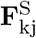 accordingly. Therefore, starting from a line configuration crowns spread along the x-axis, and perturbations are required to move along the y-axis and reach a staggered configuration. Inset: **F**^S^ as a function of crown separation *r*, normalized by *ε* (Eq. 1). **(B)** In order to extract the force scaling factor *ε* (Eq. 1) corresponding to experiments (e.g. Fig. 1), we run a parameter sweep over *ε* for simulations of 5 interacting plants, with dynamical parameters extracted from single plant experiments as detailed in the main text. We plot the final center-center distance *d*_CC_ as a function of *ε*, and find that the value *ε* = 1.29 corresponds to the experimental distance *d*_CC_ = 1.21, indicated by the dashed lines. Shaded area corresponds to two standard deviations **(C)** Example of a simulation informed by experiments, with shade avoidance scaling factor *ε* = 1.29 and step sizes sampled from the experimental distribution (Fig. 2F). Initial and final configurations of plant crowns (pink and blue circles, accordingly) over 7 simulated days, with crown centers marked by pink dots and blue triangles, respectively. Connecting lines represent crown trajectories. **(D)** The center-center distance (*d*_CC_) between pairs of adjacent crowns increases over time. The average over 12 experiments is indicated by the black line, with the shaded area representing 1 standard deviation. The red line shows *d*_CC_ over the course of the simulation in (C).

We show this approximation holds in the next section. Lastly, the range of motion of the crown is limited by the stem size [37], which in turn increases linearly in time. We therefore introduce a circular reflective boundary of radius *R*_B_(*t*) surrounding each crown in the model (depicted in Fig. 2C), and assume it increases linearly in time. Building on the force and noise terms defined above, we describe the crown center dynamics of plant i using coupled overdamped Langevin-type equations, following:

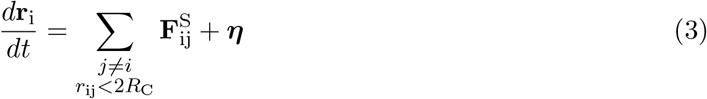

where for clarity we omit the dependence on time. All terms are informed by experiments, as detailed in the next Section.

### Model variables informed by experiments

#### Single plant dynamics

We approximate a growing plant crown as a disk whose area 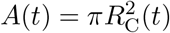 increases linearly in time with a growth rate *dA/dt*, graphically represented in Fig. 2A. We evaluate crown size from single plant experiments over 10 days. Fig. 2A shows an example of the segmentation of a plant crown, with a snapshot from the beginning of the experiment overlaid with one from the end. Fig. 2B shows *A*(*t*) as a function of time. A linear fit yields a crown growth rate of

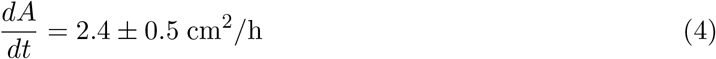

Following Eq. 3, we approximate the kinematics of single plants, driven by circumnutations, as a random walk with step sizes *η* taken from a distribution *P* (*η*) (schematically shown in Fig. 2C). The statistical characteristics of the random walk are evaluated from extracted trajectories of the crown centers from single plant experiments. An example of a trajectory spanning 10 days is shown in Fig. 2C, where discontinuities are due to the untracked movement during nighttime (8 hours). In order to quantify the circumnutation movements we first assess the sampling time step which captures the movement in a relevant way, similar to the approximation in a Langevin equation describing Brownian motion; choosing a time step which is too short over-samples the trajectory, resulting in correlated steps, and if it is too long it loses information. We therefore calculate the step-step velocity auto-correlation *C*_vv_(*t*) (defined in Eq. 9 in the Methods) as a function of different time steps, shown in Fig. 2D, and choose the shortest time step at which the auto-correlation goes to zero, at Δ*t* = 0.6 h. That is, steps Δ*t* = 0.6 h apart are uncorrelated and can be modeled as a random walk with step sizes *η* = |Δ**r**| = |**r**(*t* + Δ*t*) − **r**(*t*)|, at the basis of the noise term ***η*** in Eq. 3.

We further validate this by examining the mean squared displacement (MSD) ⟨*r*^2^ (*t*)⟩ of crown centers (given by Eq. 10 in the Methods) averaged over 8 plants each recorded over a period of up to 10 days, shown in Fig. 2E. Gaps in the MSD represents the untracked nighttime movement. The MSD agrees with a linear fit, yielding

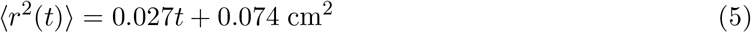

for *t* expressed in hours, with coefficient of determination *R*^2^ = 0.84. Fitting the MSD to a power-law yields a slightly sub-linear description, with a similar goodness of fit (Fig. S3 in the SM). For simplicity we adopt the linear MSD which allows us to use a regular random walk representation. Next, we extract the distribution of step sizes *P* (*η*), and find that it exhibits a remarkably broad range, spanning almost 3 orders of magnitude. The distribution is well approximated with a 2-component Gaussian mixture model (GMM), with *R*^2^ = 0.82, as well as a truncated power-law

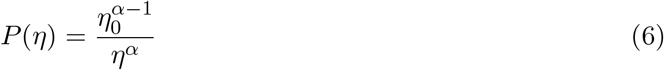

with exponent *α* = 1.40 ± 0.05 and a normalizing prefactor *η*_0_ = 6.2 × 10^*−*4^ cm (Fig. 2F). The minimal and maximal step sizes that truncate the distribution are *η*_min_ = 7.4 × 10^−4^ cm and *η*_max_ = 2.03 cm, respectively. The minimal step size is limited by the image resolution, detailed in the Methods. The normalizing prefactor, such that *P* (*η*) integrates to 1 within this range, is then 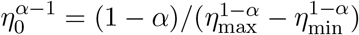. The uncertainty of the exponent is determined by bootstrapping (Methods). Although the GMM fit is slightly better, we choose to utilize the power law form as it effectively captures the breadth of the distribution using a single parameter, which proves instrumental for the analysis conducted in the following section. In Fig. S6 in the SM we show that this distribution is robust over time, and does not change significantly throughout the experiment. Lastly, we recall that the range of motion is limited by stem size, accounted for by introducing a circular reflective boundary surrounding each crown in the model (depicted in Fig. 2C). As described in the model in the previous section, the radius of the boundary *R*_B_(*t*) increases linearly in time, and without loss of generality we take it to follow the MSD in Eq. 5, such that *R*_B_(*t*) = 0.027*t*+0.074 cm^2^. In the SM we show examples of non-linear relations of *R*_B_(*t*) yielding incorrect MSD trends (Fig. S5). Put together, we model the dynamics of the fluctuating crown position of a single plant as an uncorrelated random walk with steps taken from the extracted distribution *P* (*η*) in Eq. 6, bounded by a circular reflective boundary *R*_B_(*t*) increasing linearly in time - akin to a bounded truncated Lévy flight [38, 39]. Simulations recover the trend of the experimental MSD for single plants (Fig. 2E).

#### Simulations of multiple interacting crowns recover measured self-organization dynamics

We now simulate experiments of 5 mutally shading sunflowers (as shown in Fig. 1), based on the Langevin-type model in Eq. 3, informed by experiments. Plant crowns are initially placed on a straight line 6.8 cm apart, and dynamical variables are informed from single plant dynamics characterized in the previous section, namely: the crown growth rate *A*(*t*) (Eq. 4), step size distribution *P* (*η*) (Eq. 6), and repulsive boundary *R*_B_(*t*).

To determine the scaling factor *ε* of the shading force 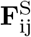 (Eq. 1), we perform a parameter sweep; running simulations over a range of values of *ε* and calculating the final *d*_CC_ for each. We find that *d*_CC_ generally increases for larger values of *ε* (Fig. 3B), thus capturing the general self-organization dynamics. The observed saturation represents the maximal deflection governed by *R*_B_(*t*): even when *ϵ* → ∞ (hard disk repulsion) the plants have nowhere to go and therefore *d*_CC_ cannot increase. We identify that *ε* ≈ 1.29 reproduces the experimental value *d*_CC_ = 1.21, as shown in Fig. 3C. Put together with the crown growth rate and the single plant perturbation dynamics, despite the anticipated experimental spread our model captures the general trends of the evolution of both the MSD of single plants (Fig. 2E), as well as the *d*_CC_(*t*) representing the self-organization in rows of multiple shading plants (Fig. 3C-D), providing a quantitative description of the system’s characteristic dynamics.

### Analysis of effects of circumnutations on the self-organization process

Having corroborated our minimal model, it now serves as a virtual laboratory, enabling us to interrogate the role of circumnutations, represented as noise, in the self-organization process, and compare to different amounts of noise. We tune the amount of noise in the system based on the spread of the step size distribution *P* (*η*). Assuming a general power-law form as in Eq. 6, the width is governed by the exponent *α*: large values of *α* correspond to narrow distributions and therefore mostly small fluctuations, while small values of *α* correspond to (truncated) heavy-tailed distributions allowing large fluctuations (illustrated in Fig. 4). In order to be able to compare across noise distributions, we require that *P* (*η*) integrates to 1 within the limiting step sizes *η*_min_ and *η*_max_ (fixed and equal to the experimental values), yielding normalizing prefactors 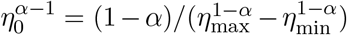. We further quantify the noise by calculating the variance, defined in Eq. 2. We substitute the general power-law form (Eq. 6) and integrate within the range [*η*_min_, *η*_max_], leading to the form

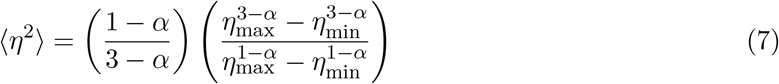

**Figure 4:**
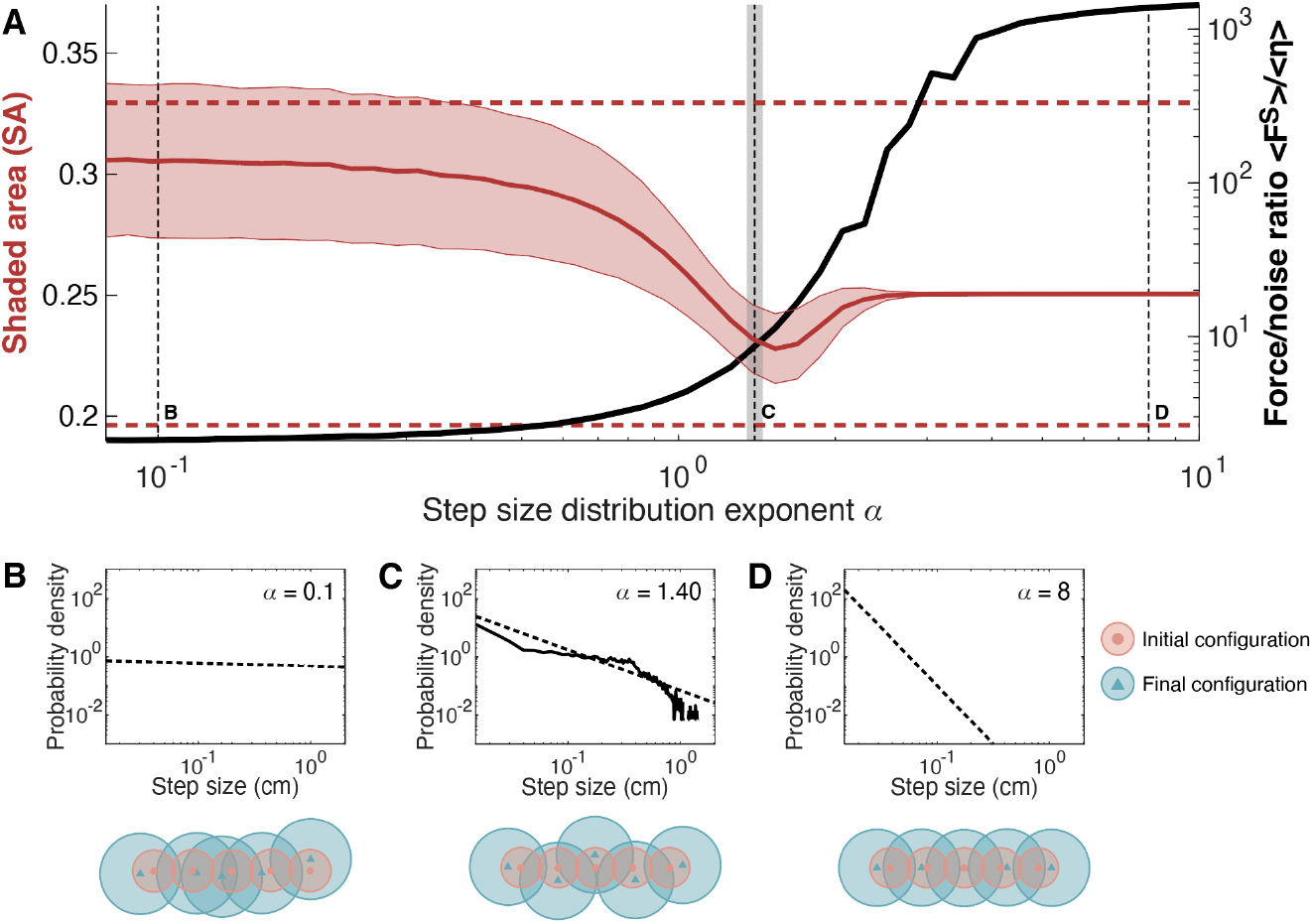
Simulations suggest circumnutations as source of functional noise. **(A) Left axis:** Fraction of shaded area (SA) as a function of *α*, the power-law exponent of the step size distribution in Eq. 6 which controls noise. The center red line represents an average over 5000 simulations, while the shaded area corresponds to two standard deviations. We identify three regimes: for high noise (small values of *α*), random movements dominate over repulsive interactions, producing disordered configurations that can result in greater shaded area. For very low noise (large values of *α*), plants remain close to their initial locations due to the lack of symmetry-breaking perturbations that push plants off the horizontal axis. In between, there is a minimum at an optimal range of *α*, where symmetry breaking results in the plants self-organizing into an alternately-deflecting “zigzag” pattern. Dashed vertical lines represent exemplary *α* values for high noise (*α* = 0.1, shown in (B)), the measured experimental value (*α* = 1.4, in (C)), and low noise (*α* = 8, in (D)). The gray shaded region represents two standard deviations about the experimental value, determined via bootstrapping. The upper horizontal red dashed line represents the shaded area if the plants do not move from their initial configurations. The lower horizontal red dashed line represents the minimum possible shaded area given the reflective boundary *R*_*B*_ on the plants’ movements. **Right axis:** The black line plots the force/noise ratio, given by the average net force on a plant at the end of the simulation ⟨**F**^*S*^⟩ divided by 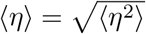 (Eq. 2). The ratio is low for small values of *α* where noise dominates, and increases with increasing *α*, saturating at large *α* where perturbations are too small to break symmetry. The minimum in SA occurs within the transition between these two regimes. **(B-D)** Dashed lines show examples of step-size distributions for *α* = 8 (low noise), *α* = 1.40 (measured value), and *α* = 0.1 (high noise), respectively, with examples of simulated final plant configurations for each respective distribution. In panel C, the experimental distribution is also shown, as a solid black line.

The dependence of noise ⟨*η*⟩ on *α* is shown in Fig. S7B in the SM; numerical values agree with the analytic form in Eq. 7, showing a sharp transition between two regimes, in line with the assumption that *α* controls noise.

We define a parameter reflecting the performance of a system of self-organizing sunflowers, in terms of the shaded area ratio (SA) of the crowns after 7 days, relative to the total crown areas (defined in Eq. 15 in the Methods): the lower the shaded area, the greater the energy the system is able to extract through photosynthesis, and the better the performance. We run simulations with parameters set by experiments, as described before, and perform a parameter sweep over *α*, which sets the noise. Fig. 4A displays the relative shaded area SA as a function of *α*, and reveals three distinct regimes. For low perturbations with *α* ≥ 2.5, the shaded area remains constant. It then decreases to a minimum for intermediate perturbations with 2.5 *< α <* 0.5, and increases monotonically for larger fluctuations with approximately *α <* 0.5. While the distributions are truncated due to physiological and resolution limitations, it is interesting to note that non-truncated power-law distributions have a well-defined mean over *η* ∈ [1, ∞) only if *α >* 2, and have a finite variance for *α >* 3. In the figure, we also highlight the shaded area for plants that remain in their initial positions over the course of growth (upper dashed line), and we observe that for some simulation instances at low *α*, the dominance of noise results in shaded area greater than this value. We also highlight the lowest possible shaded area given the reflective boundary *R*_*B*_ constraining the plants’ movements (lower dashed line), and observe that the minimum achieved for moderate *α* is close to this value, but does not reach it exactly.

Examples of final configurations from simulations corresponding to different regimes are shown in Fig. 4B-D: strong noise (*α* = 0.1), moderate noise (the experimentally measured value, *α* = 1.4), and weak noise (*α* = 8). As expected, with weak noise plants remain close to their initial linear configuration, only deflecting along the row, but not away from it. This can be understood by considering that shading neighbors on opposite sides of a crown will convey repulsive forces of equal magnitude but opposite directions, resulting in no net movement of the crown away from the line, as illustrated in Fig. 3A. As noise increases, random fluctuations cause plants to break the symmetry of the system by displacing off the horizontal axis, resulting in alternating off-axis deflections in a zigzag pattern that minimizes the shaded area. This represents the ability of the system to explore a variety of states. As noise increases further, random movements dominate over the repulsive forces, potentially leading to sub-optimal disordered configurations with higher shaded area. We observe that the experimentally-determined noise, represented by the power law exponent of *α* = 1.40 ± 0.05, is within the range of minimal shaded area, demonstrating how sunflowers may leverage circumnutations as functional noise to self-organize optimally.

To better understand these regimes, we examine the ratio of the average net force exerted on a plant 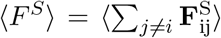 to the average noise 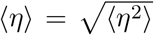 - the two terms governing the Langevin-type equation Eq. 3. This force-noise ratio represents the ratio of shading-driven motion to diffusive motion, and can be thought of as an analogy to the Péclet number. The force-noise ratio is plotted together with the shaded area SA in Fig. 4A, as a function of *α*. For low *α*, this ratio is small, due to the dominance of noise, and increases sharply for high values of *α* where the weak noise does not induce a significant out-of-row net force to break the symmetry. Comparing this to the shading area ratio SA, we find that optimal arrangements, with a minimum in SA, occur during a sharp transition between two regimes or ‘phases’, as represented by the force/noise ratio.

We note that the value of *η*_min_ is set by resolution limitations, and in reality may be smaller. In Fig. S7 in the SM we find that our results are robust to decreasing values of *η*_min_, leading to a slight shift in the position of the SA minimum closer to the experimental *α* value. Our simulations also reveal that the system is robust to variations in the repulsive interaction (Fig. S2), which may correspond to, for example, variations in the shading due to leaf thickness, distance between leaves, or fluctuations in the environmental lighting.

## Discussion

While circumnutations are ubiquitous in plant systems, and generally associated with exploratory movements, a quantitative understanding of their role is elusive. Here we report, for the first time, their role in facilitating an optimal growth pattern for a crowded group of mutually shading plants. We revisit the problem of self-organization observed for sunflowers [6], mediated by shade response interactions, suggesting circumnutations as a source of functional noise. In order to test our hypothesis, we developed a Langevin-type parsimonious model of interacting growing disks, informed by experiments, successfully capturing the characteristic dynamics of single plants as well as multiple interacting plants. This framework provided an *in silico* laboratory, enabling us to interrogate the role of circumnutations.

While traditionally circumnutation movements have been investigated in terms of the geometry of the final trajectory, here we examined the characteristic statistics of their dynamics. We found that the movements can be described as a bounded random walk characterized by a remarkably broad distribution of step sizes, or velocities, covering three orders of magnitude. This wide distribution may be related both to internal and external factors, such as ultradian and circadian rhythms, mechanical effects associated with self-weight, changes in turgor pressure due to watering, and changing light gradients, warranting further investigation. We ran simulations spanning a range of noise values *α*, and an analysis of the shading area ratio SA, describing the photosynthetic performance of the system, revealed that the experimental value occurs within a minimum in the SA, or peak performance. We then carried out an analysis of the ratio between the average net force exerted on a plant and the noise associated with circumnutations - the two terms governing the Langevin-type equation Eq. 3, and an analogy to the Péclet number of the system. We found that optimal arrangements, with a minimum in the shading area ratio SA, occur during a sharp transition between two regimes or “phases” as represented by the force/noise ratio. In all, we find that the observed breadth of the velocity distribution is beneficial, enabling the system to explore possible configurations in order to reach an optimum in terms of minimal shading, and thus solving a explore-vs-exploit problem [15, 16]. These results are also in line with the so-called “criticality hypothesis” that biological systems are tuned to phase transitions that optimize collective information processing [40, 41], enabling collective agility and responsiveness [42, 43]. We therefore interpret circumnutations as *functional* noise. Indeed, in motile animal systems such wide distributions of movement velocities are frequently identified with enhancement of behavioral processes, for example truncated power laws yielding Lévy flights, associated with animal search and foraging [44], and broad shouldered distributions related to sensory salience [17, 16]. We note that while we find here that circumnutations are beneficial, they also pose a cost to the plant, both due to the continuous change in leaf orientation (which by definition will not always be in the direction of light), as well as the mechanical cost of drooping sideways compared to growing straight. This cost-benefit trade-off needs to be addressed in future work. To the best of our knowledge this is the first report of functional noise in plant movements, and provides a theoretical backdrop for investigating how plants negotiate their environment, employing computational processes such as task-oriented processes, optimization, and active sensing.

The simple system of plants in a row serves as a minimal model for a range of ecological scenarios. On one end, our work has implications on future design of agricultural systems, as suggested by previous observations that self-organization in dense sunflower arrays give rise to an increased oil yield [6]. In tandem with advances in agronomic research and selective breeding, our results can provide insight into optimal plant spacing and harness growth dynamics to increase crop yield. On the other end, this system provides a minimal representation of natural environments such as a meadow or forest, where crowded plants shade each other in a competition for light. Understanding the dynamics of our simplified system over a wide range of timescales will be crucial to embedding the mechanics of self-organization in an ecological context. Indeed, while dynamics are considered a critical aspect of collective behavior of social organisms, this is generally overlook in plants [45]. This work provides unique insight on the role of growth dynamics on the ecological picture, a dividend of the statistical physics approach used here.

While we primarily analyze emergent self-organization in the two-dimensional top-down view of the system, we note that in fact, since plants move by growing, the system is an emergent three-dimensional structure where space and time coupled - a conceptually novel class of active matter. The shape of a shoot at a given moment in time represents its sensorial history, e.g. a plant grows towards a light source placed on its left, and when the light is switched to the right it will redirect its growth accordingly, resulting in a U-shaped stem. Therefore, neighboring plants interact not only with the current state of one another, but also their histories. An analogy can be drawn to active matter systems with memory, such as stigmergy in the social interactions of ant colonies, where ants interact with pheromone trails deposited by other ants, resulting in a collective convergence on optimal paths [46, 47]. Finally, we note that our approach can be applied to growth-driven systems other than plant organs, such as neurons, fungal hyphae, and the new generation of growing robots [48, 49, 50, 51].

## Methods

### Plant experiments

We conducted single- and multiple-plant assays using sunflowers grown from seed (‘EMEK 6’ variety, Sha’ar Ha’amakim Seeds). The seeds were first cooled in a refrigerator (seed stratification) at 5°C, peeled from their shell coats, and soaked in water for 24 hours. Each seed was then placed in a plastic test tube filled with wet Vermiculite and left to germinate in a growth chamber at 24°C, with a relative humidity of 72% and a 12:12 h light:dark photoperiod. The light intensity in the chamber was 22.05 W/m^2^.

Germination occurred after 4-7 days for each batch. Following germination, 3-to 7-cm-tall seedlings with two to four leaves that appeared healthy and well-separated were transplanted in 10 cm × 10 cm-wide and 15 cm-tall black plastic pots containing garden soil. The plants were exposed to white LED light with intensity 41.92 W/m^2^ on a 16:8 h light:dark cycle, and the setup was enclosed with black fabric to eliminate sources of reflection. The ambient temperature was approximately 26°C during the light period and 28°C during the dark period, and the humidity ranged between 43-51%. Each plant was watered with 100 mL of a 0.2% 20-20-20 NPK fertilizer solution every 2 days. Plants were maintained in this setup for approximately one week.

Plants were then selected for single-plant or multiple-plant assays that took place in conditions similar to the growth conditions described above, with the exception of light intensity at 27.05 W/m^2^. In single-plant assays (9 experiments in total), plants were maintained in the enclosed experimental setup for 7-10 consecutive days. Multiple-plant assays consisted of five plants in individual pots closely arranged side-by-side in a horizontal row, again for 7-10 consecutive days (Fig. 1A-B). Out of the total 12 multi-plant experiments 6 were on a continuous light regimen, however the characteristic statistics are similar to 16:8 light:dark regimen (Fig. S1), ruling out any effects of circadian rhythms.

### Image acquisition and tracking

We record plants during a 16-hour-long “light” period for up to 10 consecutive days. Each plant assay was recorded from a top-down view with a Logitech C270 HD webcam. Images were acquired every 5 minutes during the light period using a Raspberry Pi Model 4 single-board computer. We obtain images from 8 single-plant assays and 13 multiple-plant assays.

We perform image segmentation on videos of plants grown in individual setups using the colorThresholder function in MATLAB (MathWorks Inc., Natick, MA), from which we determine the area of the plant crown from the top-down view in single-plant assays. We segment the plant crown for every image, thus recording the crown area and the crown’s center position as a function of time.

For multiple-plant assays, plant crowns can overlap from the top-down views, complicating image segmentation. To track the movement of the crowns in both single- and multiple-plant experiments, we manually annotate the center point of each crown in the first frame of the video and track its position using the DLTdv tracking software [52].

### Plant dynamics

#### Center-to-center distances between plants

We define the position of crown i at time *t* as **r**_i_(*t*), and the initial position as 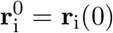. We define the average center-to-center distance between adjacent plants *d*_CC_(*t*), normalized by the initial center-center distance at *t* = 0, as

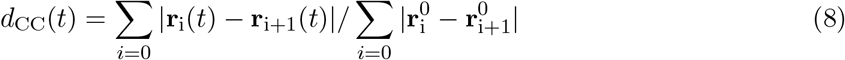

This value serves as a measure of how closely (*d*_CC_ *<* 1) or sparsely (*d*_CC_ *>* 1) the plant crowns are distributed, where *d*_CC_(0) = 1 by definition.

#### Velocity autocorrelation

To obtain a sampling rate of plant movement that captures random walk dynamics, we first quantify the autocorrelation of velocities, *C*_*vv*_(*t*). If the instantaneous velocity **v**(*t*) of the plant at time *t* is given by **v**(*t*) = (**r**(*t*) − **r**(*t* − *dt*))*/dt*, where *dt* is the length of time between subsequent frames in the recording (5 minutes), then the velocity autocorrelation as a function of time lag Δ*t, C*_*vv*_(Δ*t*), where Δ*t* = *ndt* is an integer *n* multiple of the time between frames *dt*, is given by

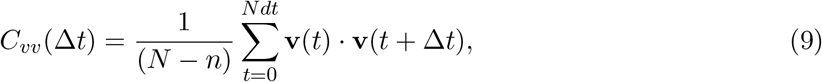

where *N* is the total number of frames in the video. We observe that *C*_*vv*_(Δ*t*) is initially positive for Δ*t <* 0.6h, before decreasing to 0 at Δ*t* ≈ 0.6 h. Hence we choose a time step in our model that corresponds to 0.6 h, such that steps taken 0.6 h apart are uncorrelated and can be represented by random walk.

#### Step size distribution

To determine the step size distribution, we compute the distribution of all steps separated by Δ*t* = 0.6h, i.e. the distribution of |*η*(*t*)| = |Δ**r**(*t*)| = |**r**(*t*) − **r**(*t* − 0.6h)| for all *t* ∈ [0, *T* ] where *T* is the length of the video. We perform a least-squares fit of this distribution to a power law, *P* (*η*) ∼ *η*^*−α*^, and find a best-fit value of *α* = −1.4. To obtain an error estimate of *α*, we perform bootstrapping by sampling from the step size distribution with replacement, fitting the resampled distribution to a power law, and repeating the resampling and fitting process for 10^4^ iterations. We then compute the standard deviation of *α* over these 10^4^ bootstrapped iterations to obtain a value of 0.05.

#### Mean squared displacement

To characterize the trajectories of plants, we compute the mean squared displacement (MSD) across all single plant recordings as

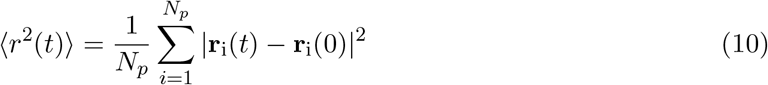

where **r**_i_(*t*) is the position of plant i at time *t*, and *N*_*p*_ is the total number of plants, in this case *N*_*p*_ = 8. There are gaps in the MSD during the nighttime periods when the plants were not recorded. We perform a least-squares linear fit of the MSD, obtaining the best-fit expression MSD(*t*) = (0.027 cm^2^/h)*t* + 0.074 cm^2^, for *t* expressed in hours, with coefficient of determination *R*^2^ = 0.84.

### Minimal model of shade avoidance response

We formulate a model of crown deflections in the 2-D plane by modeling each plant as a circular crown with an area that grows linearly in time. When the separation between a pair of crowns is less than the sum of their radii, each crown experiences a repulsive “shade avoidance” force **F**^S^ of equal magnitude (Fig. 3A).

Consider a system consisting of two crowns i and j each with radius *R*_*C*_ (we note that *R*_*C*_ implicitly depends on time, but omit this dependence for simpler notation) and separated by the vector **r**_ij_ = **r**_j_ − **r**_i_. Then, the shade avoidance force 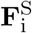 (Fig. 3A) acting on crown i is given by

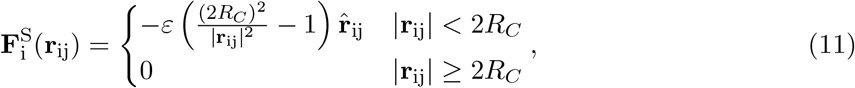

where *ε* is a coefficient that scales the magnitude of the force, 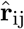 is the unit vector in the direction of **r**_ij_, and the −1 term is introduced to shift the value of the force such that it approaches 0 when |**r**_ij_| → (2*R*_*C*_)^−^ and avoids the jump discontinuity that would otherwise occur at |**r**_ij_| = 2*R*_*C*_. Furthermore, we fix 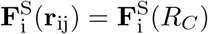 for |**r**_ij_| *< R*_*C*_ to prevent extremely large forces from occurring in the simulation.

The shade avoidance force 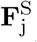 acting on crown j is of equal magnitude and points in the opposite direction along the unit vector 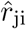. For a system of multiple mutually shading crowns, the force 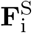 acting on crown i is the sum of all pairwise forces between crown i and all other crowns in the system.

Then, at each time step of the simulation, the change in position of crown i is given by the overdamped equation of motion

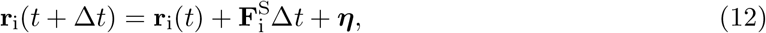

where ***η*** is a vector representing a random step, with direction sampled from the uniform distribution [0, 2*π*), and magnitude sampled from the a step size distribution. To generate the results illustrated in Fig. 3, the magnitude is sampled from the step size distribution extracted from single plant experiments (Fig. 2D). To generate the results illustrated in Fig. 4, where we vary the magnitude of the noise by changing the power law exponent *α*, we sample from a step size distribution of the form

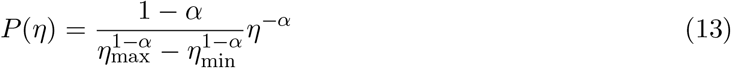

where the preceding term is a normalization factor such that *P* (*η*) integrates to 1 over the range [*η*_min_, *η*_max_]. The value *η*_max_ = 2.03 corresponds to the largest step size observed in the experiment, while the the value *η*_min_ = 7.4 × 10^*−*4^ cm corresponds to the largest distance a plant can move without being detected in a video recording (i.e., the measurement precision). The position of the plants is tracked using the DLTdv tracking software [52] (see above “Image acquisition and tracking” subsection), which interpolates positions to a precision of 0.01 pixel. Then, the largest distance a plant can move without being tracked is the 0.01 times the diagonal of one pixel, which corresponds to 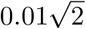 times the pixel-to-centimeter conversion rate (1/19 cm^*−*1^), or 7.4 × 10^*−*4^ cm.

Sampling from a given step size distribution is achieved using inverse transform sampling. The cumulative distribution function *CDF* (*x*) is determined from the experimental distribution; then, a random variable *u* is sampled from the uniform distribution [0, 1]. The step size magnitude is chosen to be the value *x* at which *u* = *CDF* (*x*). In the SM we verify that the variance of simulated step sizes is equal to the analytical value (Fig S7B).

Because the center of the crown is rooted to the ground via a stem of finite length and stiffness, the crown’s movement is constrained in the *x*-*y* plane by the amount by which the stem is able to deflect. To quantify this constraining region, we compute the mean squared displacement (MSD) of the crown over the course of entire single-plant experiments (Fig. 2C).

The MSD then represents the increasing radius of a circle, centered on the crown, that bounds the movement of the crown as the stem grows, allowing for greater deflection over time. The bounding circle acts as as a reflecting boundary condition: if the change in position of the crown places the crown center outside of this bounding circle, the excess movement outside the bounding circle is reflected back toward the interior of the circle.

The radius of the crown increases according to the relation

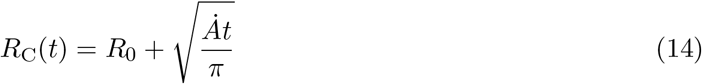

where *R*_0_ is the radius at the start of the simulation and *Ȧ* is the growth rate of the area of the crown (Eq. 4). For simplicity, in our simulations, we set the radius to be equal across each crown in the system.

We simulate multiple-plant systems with our model using experimentally-determined parameters. 5 plants are initially separated by a distance of 12.8 cm, the average value of the initial center-center distance *d*_*CC*_ without normalization. Each crown has an initial radius of 6.8 cm, which is estimated by segmenting the plant area in the first frames of multiple-plant assays, and taking this segmented area to represent five circular crowns. The area of each crown increases linearly at a rate of 57 cm^2^ (Fig. 2B) per day. Each timestep of the simulation represents 0.6 h, and the magnitudes of their random movements are drawn from the experimental step size distribution (Fig. 2E).

The system performance is evaluated by determining the fractional shaded area across all plants, given by

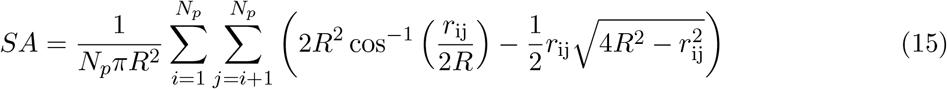

where *N*_*p*_ is the total number of plants and *r*_ij_ is the distance between the centers of plants i and j.

## Acknowledgments

O.P. acknowledges Varun Sharma, who contributed to early iterations of the computational model. O.P. and Y.M. acknowledge the hospitality of the Aspen Center for Physics, which is supported by the National Science Foundation grant PHY-1607611.

