## Supplementary Information for "Noisy circumnutations facilitate self-organized shade avoidance in sunflowers"

[1]BioFrontiers Institute, University of Colorado, Boulder, CO, USA
[2]School of Plant Science and Food Security, Tel Aviv University, Tel Aviv, Israel
[3]Zukunftskolleg, Department of Biology, University of Konstanz, Konstanz, Germany
[4]Centre for the Advanced Study of Collective Behaviour, Konstanz, Germany
[5]Department of Computer Science, University of Colorado, Boulder, CO, USA
[6]Department of Physics, University of Colorado, Boulder, CO, USA
[7]Department of Applied Math, University of Colorado, Boulder, CO, USA
[8]Department of Ecology and Evolutionary Biology, University of Colorado, Boulder, CO, USA
[9]Santa Fe Institute, Santa Fe, NM, USA

June 24, 2024

### S1. Robustness of results to nighttime

We performed a total of 12 multiple-plant experiments that lasted at approximately 7 days. In 6 of these experiments, plants were exposed to light for 15 hours a day, and were kept in the dark for the remaining "nighttime" hours; however, in the remaining 6 experiments, the plants did not experience "nighttime" and were kept under illumination for the entire experimental period. We show that this inconsistency in light conditions did not significantly affect the plants' behavior. The distributions of plants' final center-center distance $d_{CC}$ in experiments with and without nighttime are similar, as illustrated in Fig. S1. A two-sample Kolmogorov-Smirnov test does not reject the null hypothesis below a significance level of 0.05 ($p = 0.71$) that the data from the two experimental conditions are drawn from the same distribution.

### S2. Alternative shade avoidance models

We demonstrate the robustness of our results to alternative forms of the shade avoidance force used in our model. In the main paper, we use a force that scales with the squared distance $r^{-2}$ between plants (Eq. 9). We show in Fig. S2 that the shaded area as a function of noise is qualitatively similar to the result in the main paper when we use different functional forms of the shade avoidance force, namely those that scale as $r^{-1}, r^{-3}$, and $r^{-4}$, respectively.

These functional forms are as follows:

$$\mathbf{F}_{\mathrm{i}}^{\mathrm{S},1}(\mathbf{r}_{\mathrm{ij}}) = \begin{cases} -\varepsilon \left( \frac{(2R_C)}{|\mathbf{r}_{\mathrm{ij}}|} - 1 \right) \hat{r}_{\mathrm{ij}} & |\mathbf{r}_{\mathrm{ij}}| < 2R_C \\ 0 & |\mathbf{r}_{\mathrm{ij}}| \geq 2R_C \end{cases}, \tag{1}$$

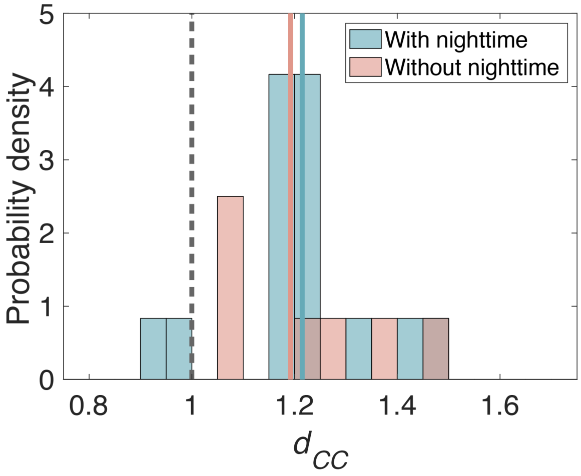


Figure S1: Center-center distances are robust to experimental light conditions. Distributions of final center-center distance between pairs of adjacent plants, $d_{CC}$, are shown for 6 experiments with a 9-hour nighttime period, and 6 experiments without nighttime (constant light exposure). A two-sample Kolmogorov-Smirnov test does not reject the null hypothesis that the data are drawn from the same distribution. Average values in both cases are shown with horizontal lines.

$$\mathbf{F}_{\mathrm{i}}^{\mathrm{S},3}(\mathbf{r}_{\mathrm{ij}}) = \begin{cases} -\varepsilon \left( \frac{(2R_C)^3}{|\mathbf{r}_{\mathrm{ij}}|^3} - 1 \right) \hat{r}_{\mathrm{ij}} & |\mathbf{r}_{\mathrm{ij}}| < 2R_C \\ 0 & |\mathbf{r}_{\mathrm{ij}}| \geq 2R_C \end{cases}, \tag{2}$$

and

$$\mathbf{F}_{\mathrm{i}}^{\mathrm{S},4}(\mathbf{r}_{\mathrm{ij}}) = \begin{cases} -\varepsilon \left( \frac{(2R_C)^4}{|\mathbf{r}_{\mathrm{ij}}|^4} - 1 \right) \hat{r}_{\mathrm{ij}} & |\mathbf{r}_{\mathrm{ij}}| < 2R_C \\ 0 & |\mathbf{r}_{\mathrm{ij}}| \geq 2R_C \end{cases}, \tag{3}$$

where the vector $\mathbf{r}_{\mathrm{ij}} = \mathbf{r}_{\mathrm{j}} - \mathbf{r}_{\mathrm{i}}$ gives the separation between two crowns i and j, both with radius $R_C$; $\varepsilon$ is a coefficient that scales the magnitude of the force; $\hat{r}_{\mathrm{ij}}$ is the unit vector in the direction of $\mathbf{r}_{\mathrm{ij}}$; and the $-1$ term is introduced to shift the value of the force such that it approaches 0 when $|\mathbf{r}_{\mathrm{ij}}| \to (2R_C)^-$ and avoids the jump discontinuity that would otherwise occur at $|\mathbf{r}_{\mathrm{ij}}| = 2R_C$. We also fix $\mathbf{F}_{\mathrm{i}}^{\mathrm{S}}(\mathbf{r}_{\mathrm{ij}}) = \mathbf{F}_{\mathrm{i}}^{\mathrm{S}}(R_C)$ for $|\mathbf{r}_{\mathrm{ij}}| < R_C$ to prevent extremely large forces from occurring in the simulation.

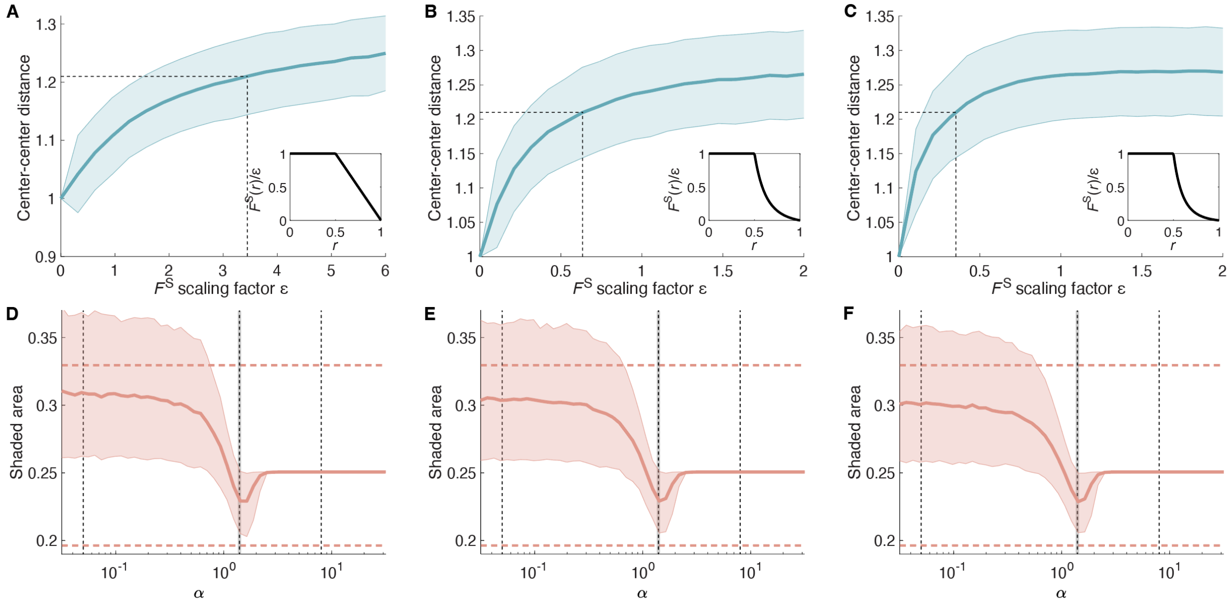


Figure S2: Results are robust to changes in the functional form of shade avoidance force. A-C: Final center-center separation $d_{CC}$ as a function of the shade avoidance force scaling factor $\varepsilon$, for shade avoidance force $F^{\mathrm{S}}(r) \sim r^{-1}, r^{-3}$, and $r^{-4}$, respectively. Dashed lines indicate the value of $\varepsilon$ that corresponds to the experimental values of $d_{CC}$. Insets illustrate these different functional forms of the force $F^{\mathrm{S}}(r)$ as a function of distance $r$. D-F: Shaded area as a function of the power law exponent $\alpha$ for shade avoidance force $F^{\mathrm{S}}(r) \sim r^{-1}, r^{-3}$, and $r^{-4}$, respectively. The force scaling factor $\varepsilon$ indicated by the dashed lines in panels A-C are used for each respective sweep. Dashed vertical line represents the experimentally-determined value of $\alpha$, while the width of the shaded gray region corresponds to 2 standard deviations as determined from bootstrapping.

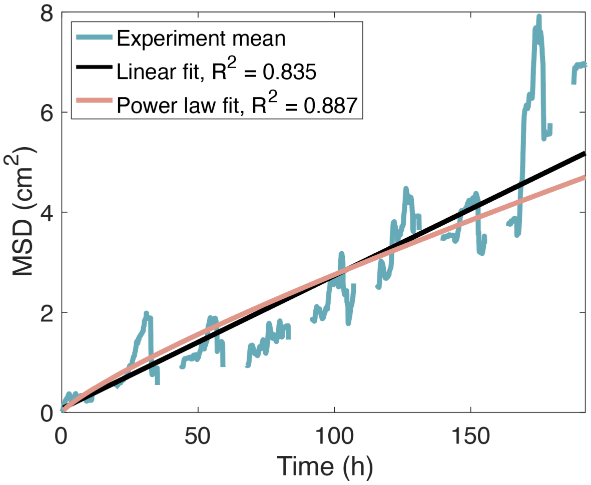


Figure S3: Comparison of linear and power-law fits to MSD. Linear fit to MSD gives MSD $= 0.027t + 0.074$. Power-law fit to MSD gives MSD $\sim t^{0.82}$.

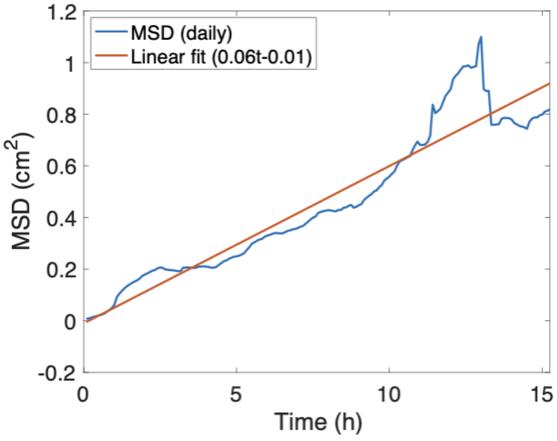


Figure S4: MSD of each day, for each experiment, averaged together, as a function of time $t$, with linear fit $0.06t - 0.01$.

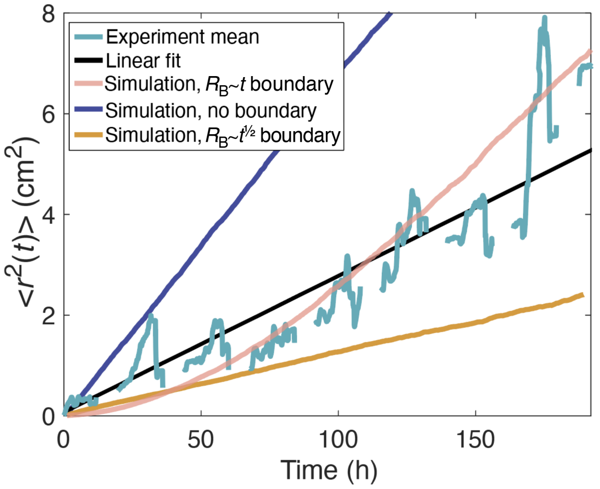


Figure S5: MSD of experiment and simulations with different boundary functions. The radius $R_{\mathrm{B}}$ of the boundary imposed in simulations described in the main text scales linearly with time $t$ as $R_{\mathrm{B}} \sim t$, and the resulting MSD (pink line) agrees with the experimental MSD (light blue line). Simulating plant movement with no boundary results in an MSD (dark blue line) that greatly exceeds that of the experimental MSD. Imposing a stricter boundary, with $R_{\mathrm{B}} \sim t^{1/2}$, results in a simulated MSD (orange line) that is much lower than the experimental MSD.

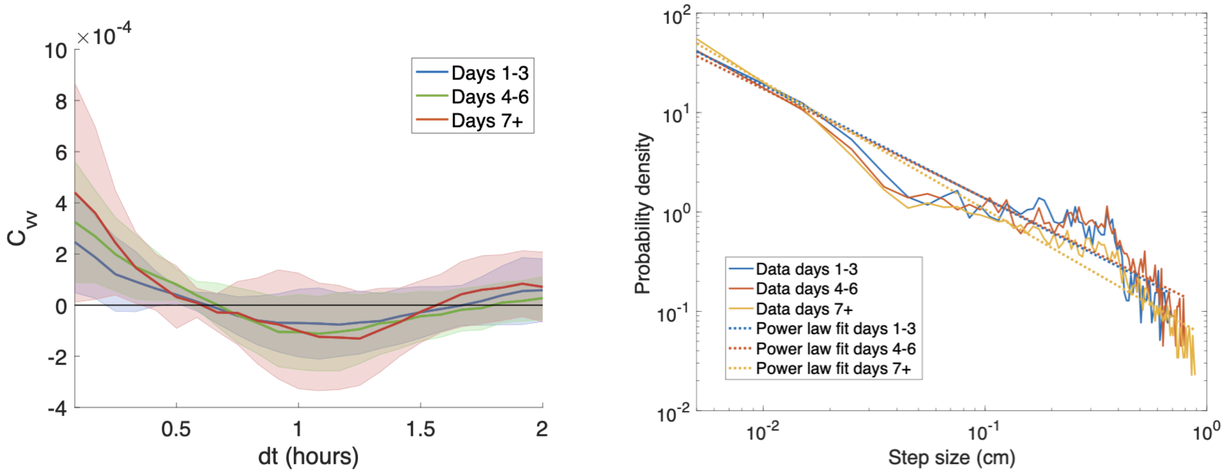


Figure S6: Autocorrelation and step size distributions split into 3 segments, where each segment corresponds to the first three days, middle three days, and final one to three days of the single plant experiments, respectively. The autocorrelation function $C_{VV}(dt)$ for each segment reaches 0 for $dt$ of approximately 0.6 h, and the step size distributions are similar across the 3 segments.

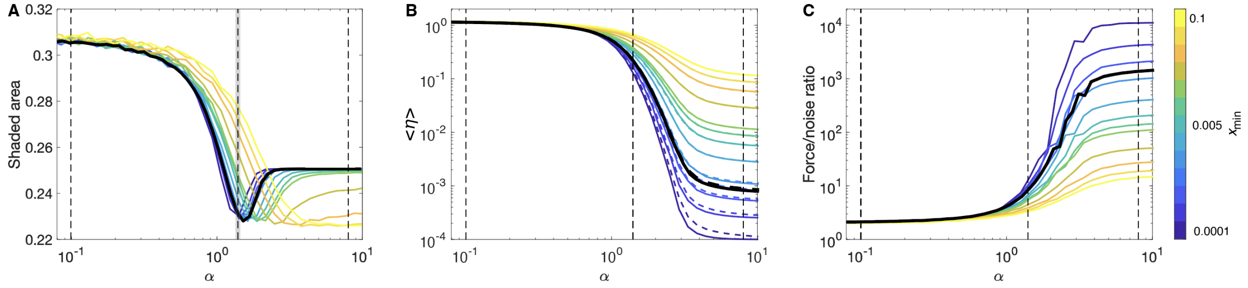


Figure S7: Shaded area ratio SA, noise $\langle \eta \rangle$, and force/noise ratio as a function of $\alpha$, for different values of minimum step size $\eta_{\min}$. Different colors represent $\eta_{\min}$ values (colorbar at far right), while the black line corresponds to the experimental $\eta_{\min}$ value of $7.4 \times 10^{-4}$. Vertical dashed lines represent $\alpha = 0.1, 1.4$, and 8, respectively, corresponding to the same values of $\alpha$ highlighted in the main text Fig. 4. A: The shaded area (SA) decreases to a minimum value as $\alpha$ increases, before rising to a constant value for high $\alpha$. The SA minimum shifts to the right as $\eta_{\min}$ increases. In the case of very large $\eta_{\min} = 0.1$, the shaded area does not again rise after reaching its minimum value. B: The magnitude of noise $\langle \eta \rangle$ as a function of $\alpha$; solid lines represent noise values measured from simulations, while dashed lines represent the theoretical values calculated using Eq. 6 in the main text. For higher values of $x_{\min}$, the simulation and theory values match almost perfectly, but these values diverge slightly for very small $x_{\min}$. When $\alpha$ is low, the magnitude of noise is high, decreasing very slowly as $\alpha$ increases. This is followed by a regime of rapidly decreasing noise when $\alpha$ takes on values that produce the SA minima seen in panel A. In the final regime, for $\alpha$ approximately greater than 4, the step size distribution is extremely narrow, with high probability assigned to small step sizes. Since the smallest possible step size is set by $\eta_{\min}$, $\langle \eta \rangle$ asymptotes to $\eta_{\min}$. C: The force/noise ratio as a function of $\alpha$. This ratio remains small, with little variation across $\eta_{\min}$ for low $\alpha$. The ratio then rises rapidly for the moderate range of $\alpha$ values in which the SA reaches a minimum, before slowly increasing for the regime of large $\alpha$ where the noise $\langle \eta \rangle$ is approximately equal to $\eta_{\min}$.

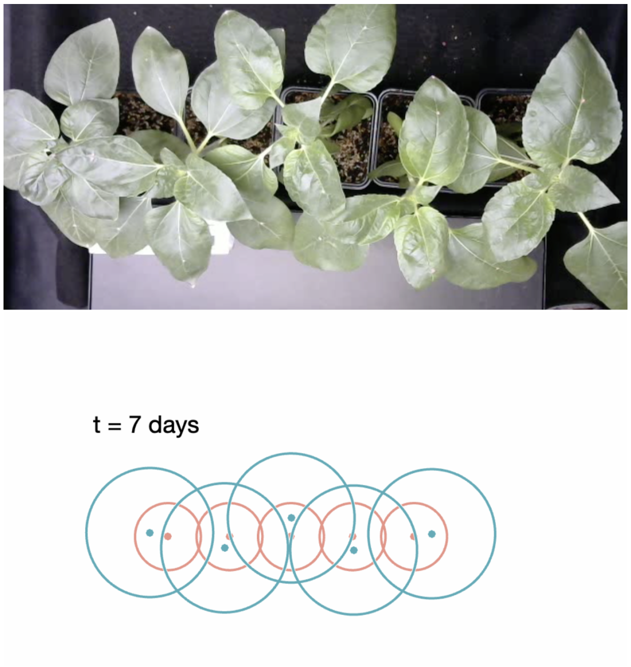


**Video S1.** Screenshot for Video 1, comparing video of multiple-plant experiment with simulation using experimentally-derived parameters.
